# Real-time audio processing of real-life soundscapes for EEG analysis: ERPs based on natural sound onsets

**DOI:** 10.1101/2021.09.20.461034

**Authors:** Daniel Hölle, Sarah Blum, Sven Kissner, Stefan Debener, Martin G. Bleichner

**Affiliations:** Neurophysiology of Everyday Life Group, Department of Psychology, University of Oldenburg, Oldenburg, Germany; Neuropsychology Lab, Department of Psychology, University of Oldenburg, Oldenburg, Germany; Cluster of Excellence Hearing4all, Germany; Institute for Hearing Technology and Audiology, Jade University of Applied Sciences, Oldenburg, Germany

## Abstract

With smartphone-based mobile electroencephalography (EEG), we can investigate sound perception beyond the lab. To understand sound perception in the real world, we need to relate naturally occurring sounds to EEG data. For this, EEG and audio information need to be synchronized precisely, only then it is possible to capture fast and transient evoked neural responses and relate them to individual sounds. We have developed Android applications (AFEx and Record-a) that allow for the concurrent acquisition of EEG data and audio features, i.e., sound onsets, average signal power (RMS) and power spectral density (PSD) on smartphone. In this paper, we evaluate these apps by computing event-related potentials (ERPs) evoked by everyday sounds. One participant listened to piano notes (played live by a pianist) and to a home-office soundscape. Timing tests showed that the temporal precision of the system is very good. We calculated ERPs to sound onsets and observed the typical P1-N1-P2 complex of auditory processing. Furthermore, we show how to relate information on loudness (RMS) and spectra (PSD) to brain activity. In future studies, we can use this system to study sound processing in everyday life.

## 1 Introduction

Mobile electroencephalography (EEG) allows to record brain activity beyond the lab while participants go about their everyday life (De Vos, Gandras, & Debener, 2014; Debener, Minow, Emkes, Gandras, & de Vos, 2012; Hölle, Meekes, & Bleichner, 2021; Wascher et al., 2021). It thereby provides unique insights into human cognition that cannot be gained by introspection or by the observation of behavior alone. Importantly, it can help us to understand the coupling of action and cognition (Gramann, Ferris, Gwin, & Makeig, 2014; Ladouce, Donaldson, Dudchenko, & Ietswaart, 2017; Parada, 2018). In everyday life, action and cognition are tightly coupled - the brain has to continuously adapt to changing events in the environment and adjust goal-directed behavior accordingly. For example, we have to react immediately to the honking of a car. By studying the brain in complex naturalistic environments, we can learn more about the dynamics of brain processes over longer time periods (Hölle et al., 2021) and in response to environmental events that are difficult to recreate in the laboratory.

If we want to understand the brain in relation to everyday life events (we will focus here on sound events), we need to capture both the brain activity and information about the events concurrently with high temporal precision: simply put, to analyze the brains’ response to the honking car at the correct moment, we need to know when the honk occurred relative to the EEG data. By relating naturally occurring sounds in everyday life to ongoing EEG activity, we can investigate questions about auditory perception and attention in real world scenarios. There is an increasing number of studies using naturalistic stimuli to study auditory perception (e.g., De Lucia, Tzovara, Bernasconi, Spierer, & Murray, 2012; Perrin et al., 2005; Roye, Jacobsen, & Schröger, 2013; Scheer, Bülthoff, & Chuang, 2018; Zuk, Teoh, & Lalor, 2020). However, to gain experimental control in this lab-based research, everyday life sounds and contexts are approximated by using artificial stimuli and conditions. With mobile EEG we can record the brain and the acoustic environment throughout the day in everyday live, and start to understand sound processing in real life contexts (Hölle et al., 2021).

Ideally, brain recordings in everyday life should not interfere with natural behavior. The used recording setup should neither be noticeable for the participant nor for outside observers, only then we can expect a natural and therefore representative behavior of the participant. Hence, the optimal setup is maximally transparent, portable, and non-restrictive for a participant (Bleichner & Debener, 2017). Such a transparent EEG solution is smartphone-based, as smartphones have sufficient computational power for EEG recordings and processing, and they can be comfortably carried in a pocket (Blum et al., 2017; Blum, Jacobsen, Bleichner, & Debener, 2019; Debener, Emkes, De Vos, & Bleichner, 2015; Hölle et al., 2021; Piñeyro Salvidegoitia et al., 2019).

To investigate sound processing in daily life, we have developed two apps for Android smartphones (Record-a and AFEx) that allow us to record and process naturally occurring sounds and brain data. Record-a is a generic app that uses the LabStreamingLayer (LSL) framework for the simultaneous acquisition and synchronization of different data streams (Blum, Hölle, Bleicher, & Debener, 2021, submitted for publication). The AFEx app dissects raw audio into acoustic features (Power spectral density, PSD; average signal power, RMS; sound onsets) in a privacy-protecting way. These features are broadcast as LSL streams and can be recorded concurrently with EEG data by the Record-a app.

Before studying sound processing in everyday life in a hypothesis-driven manner with this system, each component has to be carefully tested and validated (Scheel, Tiokhin, Isager, & Lakens, 2020). Hence, in this paper, we evaluate this smartphone-based system regarding timing precision and plausibility of the EEG data. We show how we use this system to relate EEG and acoustic features to study ERPs. We use sound onsets to calculate ERPs, spectral information to distinguish tones, and information on loudness to contrast ERPs elicited by loud and soft sounds.

## 2 Results

### 2.1 System Validation

The AFEx app needs time to buffer and process the raw audio, which creates a lag between the recorded sound onsets (onset marker) and the actual sound onsets in the real world. We conducted timing tests (see Methods) to quantify this lag and to check if it is stable over time (jitter). Table 1 shows the results of the timing tests. We identified a mean lag of 247.23 ms (62 samples at 250 Hz). When shifting the recorded sound onset markers by this number, they align with the actual onset of the sound.

**Table 1:**
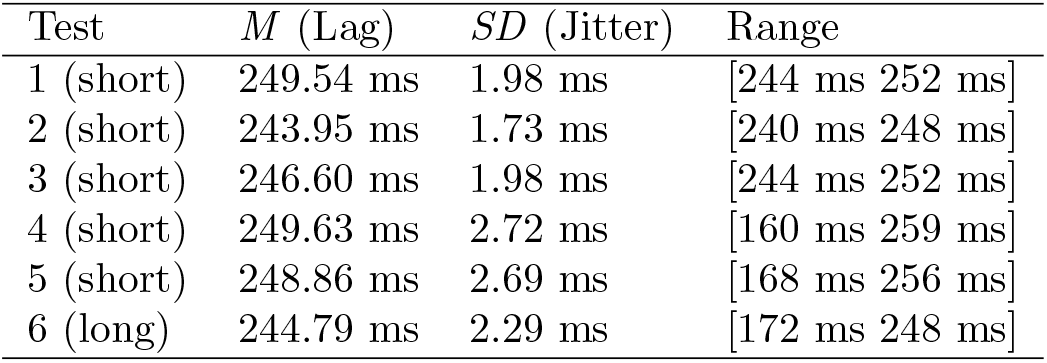
Timing Test Results. Lag describes the difference between the actual sound onset and the recorded sound onset. Jitter describes the variation of the lag over time. Short recording: 45 min; long recording: 180 min.

### 2.2 EEG recordings

Apart from timing tests, we also recorded neural data to validate the system. For the recordings, we used a smartphone running three apps: AFEx computes and streams the acoustic features; Smarting (mBrainTrain, Belgrade, Serbia), a commercial app, streams EEG data; and Record-a concurrently records the former two data streams into one file. We recorded EEG from one person equipped with a mobile EEG cap. See Figure 1 for an illustration of the system components and how they play together.

**Figure 1:**
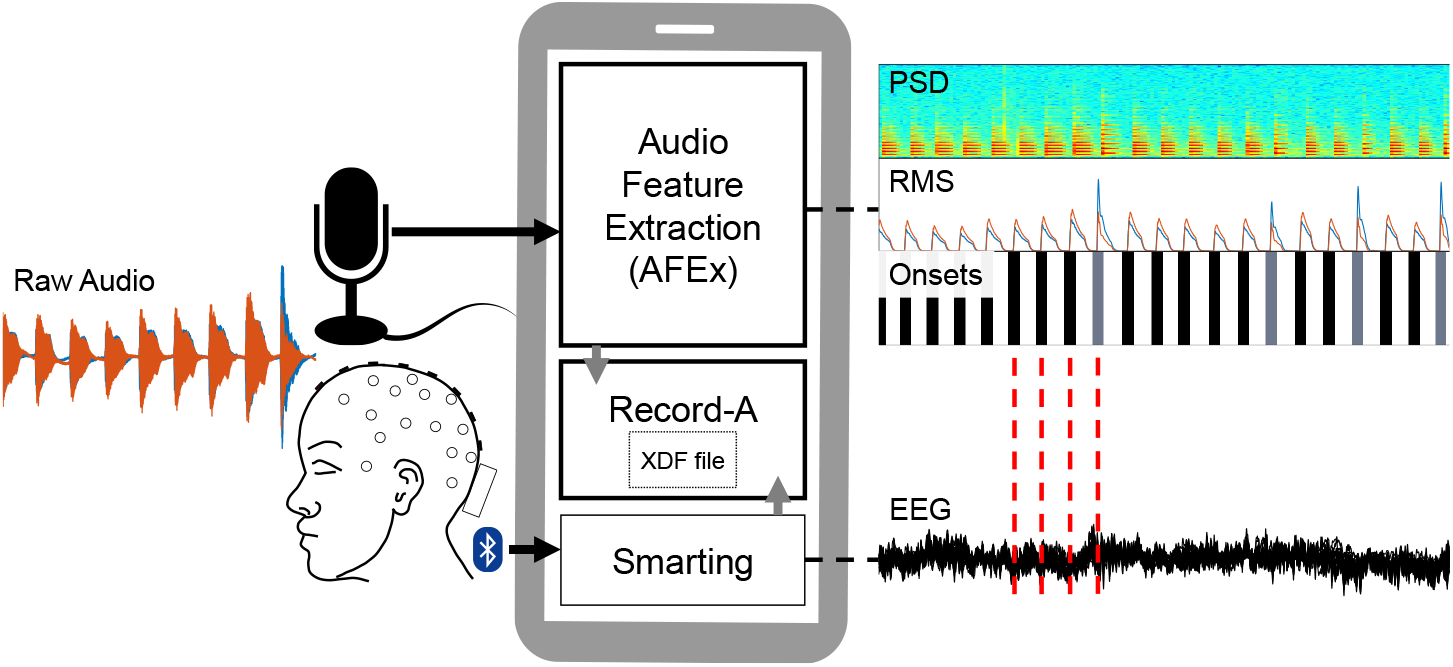
Schematic of recording setup. The participant was equipped with a mobile EEG and listened to audio. An EEG data stream was generated by the Smarting app that received the data from the amplifier via Bluetooth. The sound features (PSD, RMS, onsets) were computed by the AFEx app which received the audio data via a microphone connected to the smartphone. Audio feature and EEG data streams were recorded with the Record-a app and written in an xdf-file. With the sound onsets, we can epoch the EEG data (red lines) and compute ERPs.

The participant listened to three different auditory inputs (a random sequence of two tones, a random sequence of multiple tones, and a home-office soundscape). In figure 2, we show the average ERPs to all detected audio onsets for the three conditions. These conditions are analyzed in more detail below.

**Figure 2:**
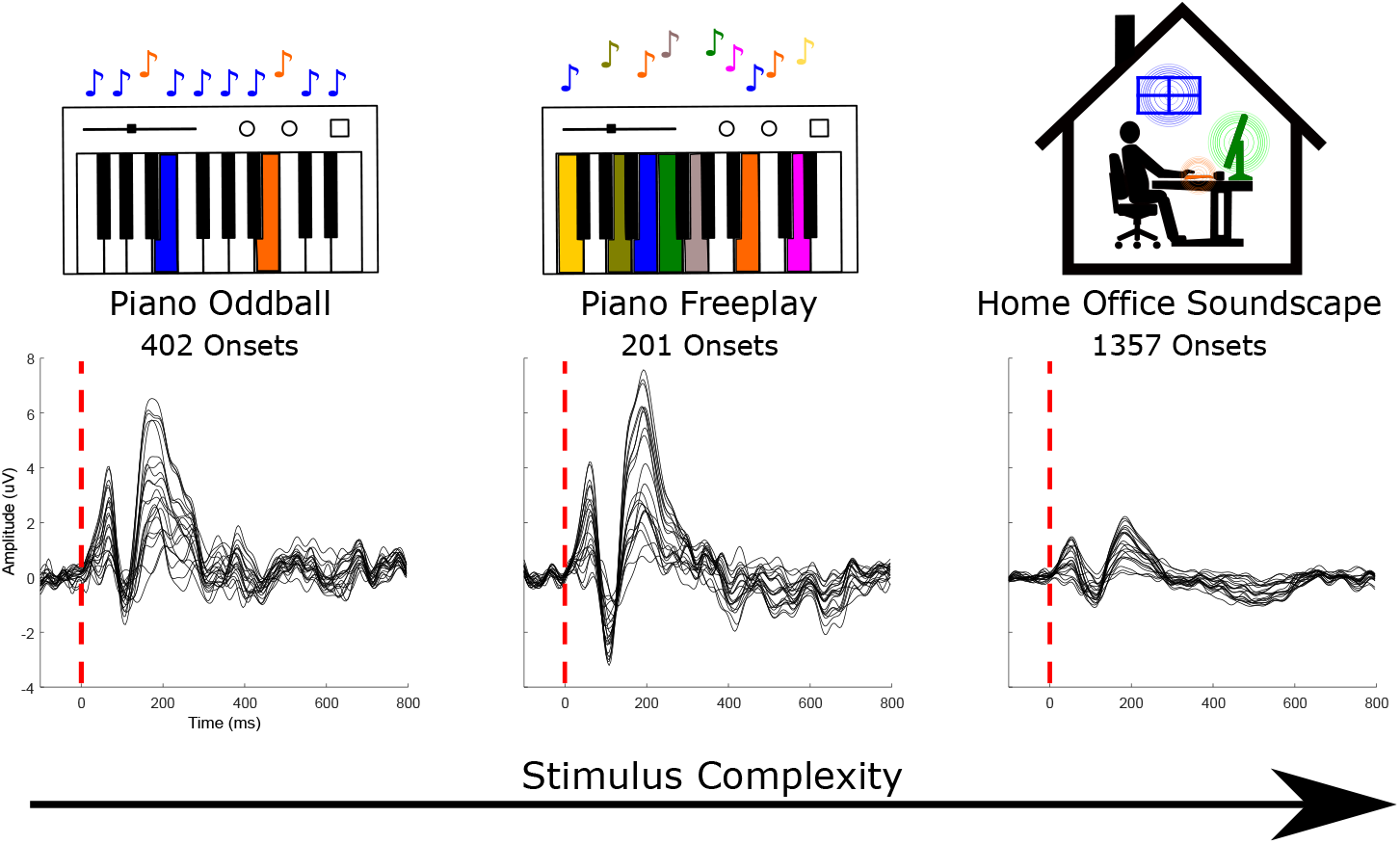
Grand average event-related potentials (ERPs) in the EEG recorded in response to the random sequence of two tones (piano oddball, 402 onsets), the random sequences of multiple tones (piano freeplay, 201) and to home-office soundscape events (1357).

### 2.3 Piano Oddball

In this condition, the participant listened to two tones (*c* and *g*) that were played live by a pianist in the same room (total duration 6 min). One note was played frequently and the other infrequently. The participant had to count the infrequent tone (target). The participant correctly counted all targets. On average, the AFEx app detected a tone every 0.99 seconds (*SD* = 0.24 s, Range = [0.25 s 1.42 s]). Figure 3a shows the results of the piano oddball, split into target and standard tones based on the PSD data. The automatic identification of targets and standards corresponded exactly to the actual notes played as recorded by a MIDI file (ground truth). For both tones, auditory evoked potentials (P1,N1,P2) can be observed. Compared to the standard tone, the target tone also shows a P3 response.

**Figure 3:**
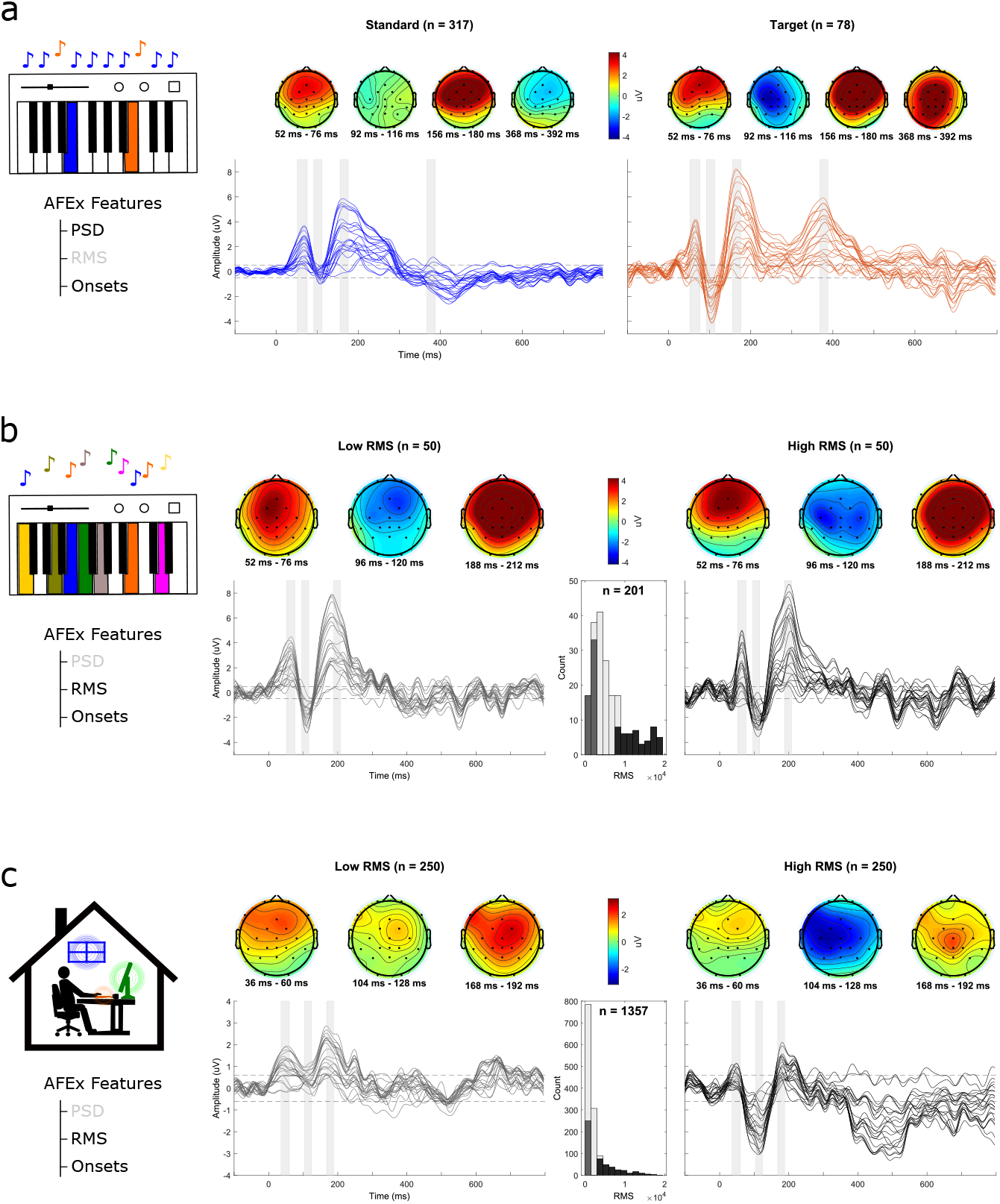
Grand average ERPs per condition. a) Average ERP to piano odd-ball, divided into standard (frequent,n=317 blue) and target (infrequent, n=78, orange) tones based on the PSD information. b) Average ERP in the piano freeplay. RMS information was used to sort audio onsets from lowest to highest RMS. The ERP is shown in response to the 50 softest (left) and 50 loudest sounds (right). The histogram shows the distribution of the RMS values (n = 201) and the color coding indicates which value range was used for the plots. c) Average ERP in response to detected sounds in the home-office soundscape (left: 250 softest sounds; right: 250 loudest sounds). The histogram shows the distribution of the RMS values (n = 1357) and the color coding indicates which value range was used for the plots. Each line represents a channel. The grey shaded arrays correspond to the time windows of the topographies. The dotted black lines represent the amplitude threshold for a signal deviation that is significantly different from baseline. Data below this threshold was considered as noise. 6

### 2.4 Piano Freeplay

The participant passively listened to the free playing of individual tones by the pianist. The pianist varied the speed and the loudness at which he played the tones. He played 329 tones (6 min) of which the AFEx app detected 201 (61.09%). On average, AFEx detected a tone every 1.82 seconds (*SD* = 1.28 s, Range = [0.39 s 9.83 s]). Figure 3b shows the average ERPs for the 50 events with the highest RMS and for the 50 events with the lowest RMS (RMS distribution: *M* = 6488.40, *SD* = 5262.06, Range = [37.17 25924.36]). Again, a P1-N1-P2 complex is clearly visible (cf. Figure 2 for the grand average of all tones).

### 2.5 Home-office Soundscape

The participant passively listened to a pre-recorded soundscape (27 min) of a home-office situation with sounds such as operating a computer and unloading a dishwasher. AFEx detected 1357 onsets. On average, it detected a sound every 1.17 seconds (*SD* = 1.52 s, Range = [0.16 s 21.49 s]). Figure 3c shows the average ERPs for the 250 events with the highest RMS and for the 250 events with the lowest RMS (RMS distribution: *M* = 2449.57, *SD* = 2746.32, Range = [805.07 21970.01]). Again, a P1-N1-P2 complex is clearly visible. A higher amplitude of the N1 component (at around 100 ms) in the high-RMS condition compared to the low-RMS one can also be observed. A paired t-test based on the mean single trial amplitudes over all channels from 104 ms to 128 ms (see topoplots) indicates that this difference is statistically significant (*t* (249) = 4.23, *p <* .001).

## 3 Discussion

We presented a system incorporating two Android apps that allow to relate acoustic features of the soundscape to continuous EEG data. Our goal was to evaluate the timing precision and EEG data plausibility of this system. In the timing tests, we showed that the temporal precision of the system is sufficient to compute ERPs. We identified a constant lag (due to signal buffering of AFEx), that can be easily accounted for by shifting the recorded onset marker, and a negligible jitter. For the EEG recordings, we presented sounds with increasing stimulus complexity from single piano tones to complex office sounds. In all experimental conditions, we found clear auditory evoked potentials in response to sound onsets. Furthermore, we showed how acoustic features in addition to sound onsets can be used to analyse EEG data. In the piano oddball condition, we used PSD to distinguish target from standard tones. The resulting ERPs show the expected P3 response for target tones only (Polich, 2007). Advancing from this simple validation task to everyday life settings, PSD information could be used to differentiate between different sound sources (e.g., Fahim, Samarasinghe, & Abhayapala, 2018). For example, in a two speaker scenario, PSD can be used to identify which speaker (low vs. high voice) is currently talking.

In the piano freeplay and home-office soundscape condition, we contrasted high-RMS sounds with low-RMS sounds. In line with literature (May & Tiitinen, 2010), we observed a significantly higher N1 response for high-RMS onsets compared to low-RMS ones in the home-office soundscape. We did not observe this difference in the piano freeplay, which may be due to the lower number of onsets and thus lower variance of RMS values. Note that ERP amplitudes in the home-office condition were lower than in the other conditions, which is most visible in the grand average of all sound onsets. This finding can be explained by the higher number of low-RMS onsets (see histogram) in this recording, resulting in lower amplitudes overall.

To calculate these ERPs, we relied on sound onsets. Importantly, there are many different methods of defining sound onsets (e.g.; Böck, Krebs, & Schedl, 2012; Thoshkahna & Ramakrishnan, 2008); for example, some methods are specifically geared for detecting onsets in music (Bello et al., 2005; Haumann et al., 2021). For the online detection of onsets, we optimized the parameters to detect clear, isolated sound onsets in the presence of background noise. Consequently, not all onsets that that humans perceive as onsets are picked up by the detector. For example, in the freeplay condition, 120 notes were missed by the detector. Thus, the onsets that were detected are a subset of all onsets a person perceives. Depending on the use case, the parameter settings can be adapted and optimized for different sound onsets.

Obviously, the acoustic features that are provided by AFEx only provide limited information about the soundscape. The app does not provide information about many aspects that are known to be relevant for sound perception; for example, whether a sound was self-generated or generated by another sound source (e.g., Sanmiguel, Todd, & Schröger, 2013), whether the sound was relevant or irrelevant (e.g., Dehais, Roy, & Scannella, 2019; Holtze, Jaeger, Debener, Adiloğlu, & Mirkovic, 2021; Scheer et al., 2018), or expected or unexpected for a person (e.g., Dalton & Fraenkel, 2012; Koreimann, Gula, & Vitouch, 2014). How to assess these factors remains a challenge. Some of these factors could be assessed by asking the person about subjective experience of the soundscape on a regular basis using momentary ambulatory assessment (Trull & Ebner-Priemer, 2013), but this will interrupt the person in their normal activity. An alternative is to use additional microphones that provide more information about the origin of a sound, or sensors that provide context information. However, it is unlikely that we will be able to capture the full complexity of the participants environment in the near future.

To sum up, we demonstrated that the presented technical setup reliably combines EEG and acoustic features and thereby allows to measure event related potential in response to everyday sounds. This smartphone-based setup can be used fast and easily. It allows us to study brain activity everywhere: We could record EEG next to a construction site, while driving through the city, or in the classroom. These contexts are already of interest to researchers (e.g., Getzmann, Arnau, Karthaus, Reiser, & Wascher, 2018; Ke, Du, & Luo, 2021; Ko, Komarov, Hairston, Jung, & Lin, 2017). In these everyday contexts, we can also investigate individual differences in sound processing: Why do some individuals notice some sounds that other people miss? How does their brain response to sounds differ? Which sounds are distracting or even annoying to an office worker on the neural level? As evident from these various examples, the AFEx app in combination with Record-a and a commercial EEG app facilitate new research of auditory processing in everyday life, which was henceforth typically only assessed with surveys (e.g., Banbury & Berry, 2005; Oseland & Hodsman, 2018).

## 4 Materials and Methods

### 4.1 Software

The Smarting app provided EEG data as LSL stream, the AFEx app computed acoustic features (sound onsets, PSD, RMS) and provided them as LSL streams, and the Record-a app recorded these LSL streams into one xdf-file. In our analyses, we could use the audio onsets provided by AFEx to cut the continuous EEG data and average over these epochs to gain ERPs (see Figure 1).

#### 4.1.1 Smarting

For streaming EEG data, we used a Smarting (mBrainTrain, Belgrade, Serbia) system. The ongoing EEG signal was provided as an LSL stream by the Smarting android app (Version 1.6.0). Note that any EEG system that provides the EEG data as LSL stream can be used.

#### 4.1.2 AFEx

AFEx is an Android app that calculates acoustic features (https://doi.org/10.5281/zenodo.5128051). It implements a framework that allows to concatenate various stages such as audio capture, signal filtering, and signal transformation to produce the desired metrics. The produced data is pushed to the LSL in chunks equivalent to the block sizes specified below. Fig. 4 shows the signal path.

**Figure 4:**
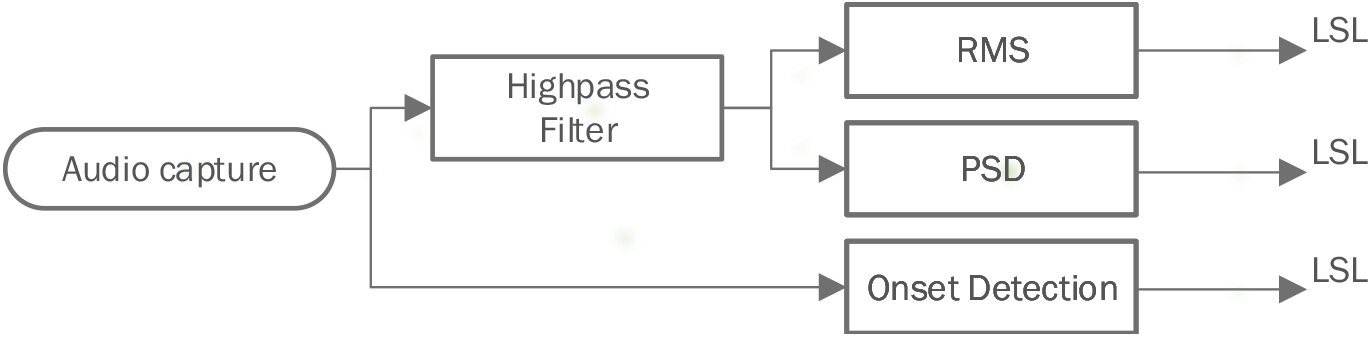
Signal path through the different stages of the Audio Feature Extraction Framework AFEx.

The different metrics are calculated and parameterized as follows:

**Audio capture** A stereo microphone signal is captured at a sampling rate of 16 kHz, buffered and transferred to subsequent stages in chunks of 250 ms. Each stage is rebuffering data to provide the blocksizes (incl. overlaps) requested by succeeding processing stages. Note that the properties of the signal are depending on the recording setup. A signal being recorded using an Android’s device internal microphone or input via the headphone/mic jack will usually result in two identical, i.e. duplicated, mono channels. Stereo signals can be recorded using external USB devices.

**Highpass filter** The second order biquad filter has a cutoff frequency of 250 Hz to suppress low frequency noise. The signal is filtered in blocks of 25 ms.

**Signal power (RMS)** The RMS is calculated with a blocklength of 25 ms and an overlap of 50 %, resulting in a sampling rate of 80 Hz.

**Spectra (PSD)** The power spectral density is calculated for blocks of 25 ms with an overlap of 50 %, resulting in a sampling rate of 8 Hz. The resulting spectra are exponentially smoothed and averaged, with one spectrum being produced every 125 ms. This method prevents reconstruction of the time domain signal to protect a person’s privacy.

**Acoustic onset detection** To detect acoustic onsets, a variable state filter produces low-, band-, and highpass filtered signals 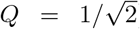. Those are then smoothed with both, slow and fast time constants, tuned to different values per band (Table 2). If the faster signal emerges from the slower in any band, an onset is triggered.

**Table 2:** Smoothing time constants for the low-, band-, and highpass-filtered signals used for acoustic onset detection.

|  | lowpass | bandpass | highpass |
| --- | --- | --- | --- |
| slow | 160 ms | 20 ms | 1 ms |
| fast | 4 ms | 2 ms | 2 ms |

All calculations are performed online on the smartphone and the results are written to LSL. The audio signal does not need to be stored or cached on the device’s permanent storage. Additionally, the parameters have been chosen in such a way as to prevent reconstruction of semantic content or information, i.e., from PSDs, thus ensuring a participant’s privacy (Bitzer, Kissner, & Holube, 2016; Kissner, Holube, Bitzer, Technology, & Sciences, 2015). Note that modified parameters or new features would require a re-validation of said privacy.

AFEx has been tested with Android OS up to version 10. Permissions to access storage and the microphone are explicitly required. Processing runs as foreground service, giving it some precedency over other apps. There are, however, no additional mechanisms in place to prevent interaction with the app, provide some sort of kiosk mode, or notify about system events like low-battery.

#### 4.1.3 Record-a

The Record-a app allows to record LSL data streams in the network into one xdf-file. With the LSL framework, different data streams with different sampling rates can be recorded simultaneously. Each data stream has its own time information in reference to a master clock and upon import, they are time corrected and thereby synchronized. This app is described in detail in Blum et al. (2021, submitted for publication). The app is available at https://github.com/s4rify/Pocketable-Labs.

### 4.2 Hardware

#### 4.2.1 EEG System

We used a 24-channel cap (easycap GmbH, Germany) connected to a mobile 24-channel Smarting mobi amplifier with 24 bit resolution (mBrainTrain, Belgrade, Serbia). We recorded the EEG data with a sampling rate of 250 Hz. The amplifier transmitted the data to the smartphone via bluetooth.

#### 4.2.2 Smartphone and Microphone

We used a Google Pixel 3a smartphone (Android Version: 10). To record stereo sound with this phone, we used customized ear microphones comprising two omni-directional EK-23024 microphones (Knowles Electronics, Illinois, USA) placed into behind-the-ear shells. The ear-microphones were connected to the phone via the USB-C port with a sound adapter (Andrea Pure Audio USB-MA, Andrea Electronics, Bohemia, USA). Naturally, the microphones process sounds differently than human ears: they may pick up on sounds not perceptible to human ears, but they may also miss sounds that the ear would perceive. However, the microphone clearly detects the onset sounds that are essential to our analyses.

### 4.3 System Validation

To relate EEG data and audio onsets reliably, both data streams need to be stable over time and have sufficient temporal precision in the millisecond domain. As the audio signal is buffered and then processed, there is a lag between the actual audio onset and the recorded audio onsets (onset marker) as they occurred in the real world. In the system validation, we conducted timing tests to quantify this lag. Furthermore, we also checked whether this lag remains stable over time (jitter).

We conducted five short (∼ 45 min) and one long (∼180min) timing tests. The smartphone was restarted between timing tests. For these timing tests we used wav-files that were generated with Matlab (Version: 9.6.0.1335978; The Mathworks Inc., Natick, MA, USA). The wav-files contained square wave impulses that were presented at 1 sec intervals at a sampling rate of 44100 Hz. We used square waves as they produce a sharp response in the EEG amplifier. The square wave impulse had a duration of 50 ms. For the short timing test, the wav-file consisted of a 10 s silence at the beginning followed by 2500 impulses. The wav-file of the long-term timing test consisted of 4 times the wav-file for the short timing test for a total of 10000 impulses. These sound files were presented via PC on external speakers (Wavemaster 1520, Bremen, Germany) in a soundproof cabin and simultaneously fed into the amplifier via a customized adapter.

Hence, with this procedure, an impulse produced both a response in the data recorded with the EEG amplifier and an audio onset as detected by the AFEx app. We can then cut the sound signal recorded with the EEG amplifier based on the audio onsets and calculate lag and jitter. The AFEx app detected onsets for all impulses plus a few extra sounds (experimenter leaving the room, nearby construction work). In our analysis (also in Matlab), we deleted these additional onsets.

We then epoched the data from −1 s to 0.1 s (baseline correction −1 to - 0.9) relative to the detected sound onsets. To quantify the delay between audio onset and onset marker, we identified the time point when the audio onset in the EEG reached its half maximum (same time point for all recordings) and then calculated the time difference between this maximum and the onset marker for every epoch.

### 4.4Participant

We recorded one volunteer (male, age between 25 and 35, ambidextrous) who had previous experience with EEG recordings. He provided written informed consent prior to participation. We deemed neural data from one participant sufficient to demonstrate the functionality of the technical setup.

### 4.5 Procedure

For the EEG recordings we had three conditions that increase in stimulus complexity. In the first two conditions, the participant listened to piano notes played live by a pianist. In the third condition, the participant listened to an home-office soundscape. In figure 2, we show the average ERPs to all detected audio onsets for the three conditions.

Before the participant was fit with the EEG cap, his skin was cleaned with alcohol and electrolyte gel (Abralyt HiCl, easycap GmbH, Germany). All impedance was kept below 10 kOhm. The ear microphones were plugged into the smartphone and placed on the participant’s ear. The smartphone was always placed on a table adjacent to the participant. For condition one and two, the participant sat in a chair approximately at a distance of 1 m to the piano (Yahama Digital Piano P-35, Hamamatsu, Japan; set to 75 percent volume), fixating his gaze on a fixation cross on a wall. The participants sat with his back to the piano; hence, he could neither see the pianist nor the experimenter. In the third condition, the participant sat in a soundproof cabin looking at a fixation cross on a computer screen while listening to a soundscape that was presented free-field (two Sirocco S30 loudspeakers, Cambridge Audio, London, United Kingdom) at a participant-comfortable volume. For all conditions, the participant was instructed to move as little as possible.

#### 4.5.1 Piano Oddball

The participant listened to live played notes on the piano. The pianist either played a *c* (159 in total, standards) or a *g* (39 in total, targets) on the keyboard. The sequence of tones was predefined and provided on a paper sheet. We recorded this oddball twice with a different predefined sequence in the second run. The participants task was to count the target tones. To check whether the pianist played exactly the notes from the sheet, we also recorded the MIDI file of the oddball. Each oddball took 3 minutes. For later analysis, we merged both oddball datasets.

#### 4.5.2 Piano Freeplay

The participant listened to the pianist playing a monophonic sequence of random tones with variable loudness and tempo for 6 minutes. The exact sequence of sounds was recorded as a MIDI file.

#### 4.5.3 Home-office Soundscape

In this condition, the participant listened to a soundscape of a home-office environment for 27 minutes. This soundscape (wav-file) included typical home-office activities such as operating a computer (clicking with the mouse, typing on the keyboard, notification sounds), walking around, using a hole puncher, or unloading the dishwasher. It did not contain music or speech. Due to technical interference in this recording, we recorded the sound onsets again in an empty room and mapped them onto the EEG recording.

### 4.6 Data Analysis

Data were analysed offline with Matlab (Version: 9.6.0.1335978; The Mathworks Inc., Natick, MA, USA) and EEGLAB (Version: v2019.0; Delorme & Makeig, 2004) using custom scripts (available at https://osf.io/bcfm3/). Filters were used as implemented in EEGLAB (zero-phase Hamming windowed sync finite impulse response).

#### 4.6.1 Preprocessing

The EEG data were re-referenced to linked mastoids, low-pass filtered at 25 Hz (Order 134) and high-pass filtered at 0.1 Hz (Order 8250). To account for the previously determined time lag, the recorded markers for sound onsets were shifted by −248ms (−62 samples; see system validation results). We deleted some markers at the beginning and the end of each recording (14 in total) as these onsets were due to the starting and stopping of the recording. We used artifact subspace reconstruction (ASR) to clean the data. For ASR, we used the EEGLAB-plugin *clean rawdata* (Version: 1.0; parameters: flatline criterion = 60, high-pass = [0.25 0.75], channel criterion = off, line noise criterion = off, burst criterion = 20, window criterion = off). We epoched the data from −0.1 s to 0.8 s relative to each sound onset (baseline correction −0.1 s to 0 s).

##### Piano Oddbal

Based on the recorded PSD data, we identified target and standard tones. First, we visually identified the first target tone in the spectrum. Second, we created a template based on this target tone. We then correlated this template with every tone in the spectrum. This procedure allowed us to automatically identify target tones when the correlation was higher than 0.9.

##### Piano Freeplay and Home-office Soundscap

In both conditions, for each audio onset, we computed an average RMS based on the mean of five samples (62.5 ms) after the audio onset of both audio channels to gain a corresponding RMS value for each sound onset.

#### 4.6.2 Random events

To obtain an estimation for the reliability of the ERPs, we calculated an amplitude threshold based on ERPs extracted from random events for each condition. Data within this threshold can be considered as noise. For 1000 permutations, we inserted 500 markers at random time points in the data, epoched with these markers, and computed the ERP over all channels. We then determined the maximum and minimum amplitude of the mean over all permutations plus two times the standard deviation. This value corresponds to approximately 95% of the amplitude distribution.

## 5 Open Practices Statement

The EEG datasets are available from the corresponding author on request. The code used for the analyses, the stimuli used, and the datasets from the timing test are available at https://osf.io/bcfm3/. The code for the AFEx can be found at https://doi.org/10.5281/zenodo.5128051. The Record-a app can be found at https://github.com/s4rify/Pocketable-Labs.

## 6 Author Contribution

DH and MB conceived the experiments with the help of SB and SD. DH carried out the data collection and analyses. DH wrote the manuscript with the input of MB, SD, SB and SK. SB lead the development of the Record-a app. MB designed the AFEx functionality and SK developed the AFEx app.

## 7 Acknowledgement

We thank Arnd Meiser for the fruitful discussions. We thank Inga Holube and Jörg Bitzer for their help in starting the development of the AFEx app. We thank the cluster of excellence “Hearing4All” for some seed funding from the “innovation pitch” competition. This work was funded by the Deutsche Forschungsgemeinschaft (DFG, German Research Foundation) under the Emmy-Noether program - BL 1591/1-1 - Project ID 411333557.

## 8 Competing Interests Statement

The authors have no conflict of interests to declare.

## Notes

https://osf.io/bcfm3/

### Competing Interest Statement

The authors have declared no competing interest.

